# Sensing the heart, seeing the world: Neural mechanisms of attentional allocation between interoception and exteroception

**DOI:** 10.64898/2026.09.22.751480

**Authors:** Sukhbinder Kumar, Joel S Winston, Hugo D Critchley, Timothy D Griffiths

## Abstract

Adaptive behaviour requires flexible allocation of attention between sensory information arising from the external environment (exteroception) and signals originating within the body (interoception). While the neural mechanisms supporting exteroceptive attention are well characterised, considerably less is known about how attention is allocated between competing interoceptive and exteroceptive information. We used functional magnetic resonance imaging to compare cardiac interoceptive attention during heartbeat counting with a matched visual target counting task. External visual input was identical during both tasks, and behavioural accuracy and subjective confidence did not differ significantly between conditions, providing a controlled comparison in which attention was directed towards internal or external information. Cardiac interoceptive and visual exteroceptive attention engaged largely distinct cortical networks. Cardiac attention recruited a distributed insula-centred network supporting interoceptive processing and cognitive control, whereas visual attention engaged a dorsal frontoparietal network and selectively shaped visual cortical activity, enhancing regions representing task-relevant central visual information while suppressing regions representing task-irrelevant peripheral information. Despite this functional segregation, activity within these systems depended on attentional allocation. Exteroceptive attention suppressed posterior insula activity, suggesting down-regulation of interoceptive processing, whereas peripheral visual regions remained suppressed during interoceptive attention, suggesting down-regulation of externally directed sensory processing. More accurate heartbeat counting was associated with less engagement of the right inferior parietal lobule, consistent with reduced bottom-up attentional capture by external sensory information. Together, these findings support competitive allocation of attentional resources between neural systems supporting internal bodily signals and external sensory information, providing a system-level account of interoceptive and exteroceptive attention.

## Introduction

Interoception, the perception of the internal physiological state of the body, has emerged as a fundamental component of human cognition, shaping emotion, self-awareness, decision-making, and homeostatic regulation (Craig, 2002; Critchley and Harrison, 2013; Chen et al., 2021; Wang and Chang, 2024). Interoceptive processing largely operates unconsciously, as the afferent axis of homeostatic and allostatic regulation. However, conscious access to internal bodily sensations also underpins motivational and affective feelings, plausibly triggering and enhancing the experience of emotions (Quadt et al., 2018). Among the various interoceptive modalities, cardiac interoception has been the most extensively studied in humans because the heartbeat provides a series of discrete, objectively measurable physiological events that are continuously available and largely outside voluntary control. Using behavioral paradigms such as heartbeat counting (Schandry, 1981) and heartbeat discrimination (Whitehead et al., 1977) or simply directing attention towards cardiac sensations, functional neuroimaging studies have identified a distributed insula-centered network supporting cardiac interoception, including the posterior and anterior insula (Critchley et al., 2004; Pollatos et al., 2007; Simmons et al., 2013; Wiebking et al., 2014b; Wiebking et al., 2014a; Wiebking et al., 2015; Tan et al., 2018; Failla et al., 2020). These findings have substantially advanced our understanding of how internal bodily signals are represented within the brain. In everyday behavior, however, internal bodily signals compete with sensory information arising from the external environment for access to limited attentional resources. Successful behavior therefore depends on the ability to flexibly prioritize either internal bodily signals or external sensory information according to current behavioral goals.

The neural mechanisms underlying such flexible prioritization have been investigated extensively, primarily within the domain of exteroceptive attention. Influential models of selective attention propose that, because processing capacity is limited, simultaneously available sources of information compete for neural representation and behavioral control. This competition is resolved through the enhancement of behaviorally relevant information and the suppression of competing inputs, thereby enabling flexible goal-directed behavior (Desimone and Duncan, 1995; Kastner and Ungerleider, 2000; Reynolds and Chelazzi, 2004; Carrasco, 2011). Consistent with these theoretical models, behavioral, electrophysiological and neuroimaging studies have demonstrated that directing attention towards one source of sensory information enhances processing of task-relevant inputs while attenuating responses to competing information (Mozolic et al., 2008; Keitel et al., 2013). However, whether these principles extend to competition between internally generated bodily signals and externally derived sensory information remains largely unknown.

Previous neuroimaging studies have largely focused on identifying the neural basis of interoceptive attention rather than examining how attention is allocated between internal bodily signals and competing external sensory information. An important next step is therefore to compare interoceptive and exteroceptive attention under conditions in which sensory input and task demands are closely controlled, allowing the neural mechanisms supporting the allocation of attention between internal and external sources of information to be examined.

To address this question, we compared cardiac interoceptive attention, measured using the heartbeat counting task, with a carefully matched visual target counting task during functional magnetic resonance imaging (fMRI). Crucially, identical external visual input was maintained during the active periods of both tasks, while attention was selectively directed towards either internal cardiac signals or external visual targets. Furthermore, behavioural accuracy and subjective confidence did not differ significantly between the two tasks, making differences in task difficulty or performance unlikely explanations for the observed differences in neural activity. This design therefore provided a controlled framework for investigating how the brain allocates attention between competing internal bodily signals and external sensory information.

The present study addressed two related questions. First, we examined whether cardiac interoceptive and visual exteroceptive attention engage distinct or overlapping neural systems. Second, we asked whether directing attention towards one source of information is accompanied by suppression of neural activity associated with the competing source. By dissociating attentional allocation from differences in sensory input and behavioral performance, the present study provides a systems-level investigation of how the brain balances attention between the internal bodily milieu and the external sensory environment.

## Methods

### Participants

Twenty healthy adults participated in the study (age range: 21–54 years; mean age = 34.3 years; SD = 9.8 years; 11 females). All participants had normal or corrected-to-normal vision. Participants who routinely wore corrective glasses were provided with MRI-compatible corrective glasses during scanning. All participants provided written informed consent before the experiment and received financial compensation for their participation. The study protocol was approved by the University College London Research Ethics Committee.

### Stimuli

Participants were scanned in a supine position inside the MRI scanner. They viewed the visual stimuli through a small mirror mounted on the scanner head coil. The mirror reflected an image cast by a projector onto a rear-projection surface placed at the back of the scanner bore. The viewing distance was 57 cm, which covered 4 degrees of visual angle. Within this viewing area, visual targets were small, striped Gabor patches spanning a visual angle of 0.083 degrees, tilted at 45 degrees to the right with a frequency of 2 cycles per degree. These visual targets were embedded within a fast-moving, 10 Hz visual stream, which also consisted of noise and distractor patches. The root-mean-square level of the noise (0.12) was adjusted based on a pilot experiment with the objective of making the visual and heartbeat counting tasks equally difficult with similar accuracy levels. Distractor patches were identical to the targets except they were tilted 45 degrees to the left. Because the visual targets were exceptionally small and presented centrally, this tight spatial constraint forced participants to focus their top-down spatial attention entirely on the center of the display to detect them. Visual stimuli were generated using custom-code based on MATLAB/Cogent software.

### Experimental Paradigm

The experiment employed a block design comprising 18 task blocks, with nine blocks each of heartbeat counting (HBC) and visual target counting (VTC). Each block began with a 3 s visual instruction cue (“Count your heartbeats” or “Count visual targets”), informing participants whether to direct their attention towards their internal cardiac sensations or towards the visual stimuli (Figure 1). The instruction cue was followed by a task period lasting 15, 20, or 25 s, with block durations pseudorandomized across the experiment to minimize temporal expectancy.

**Figure 1.**
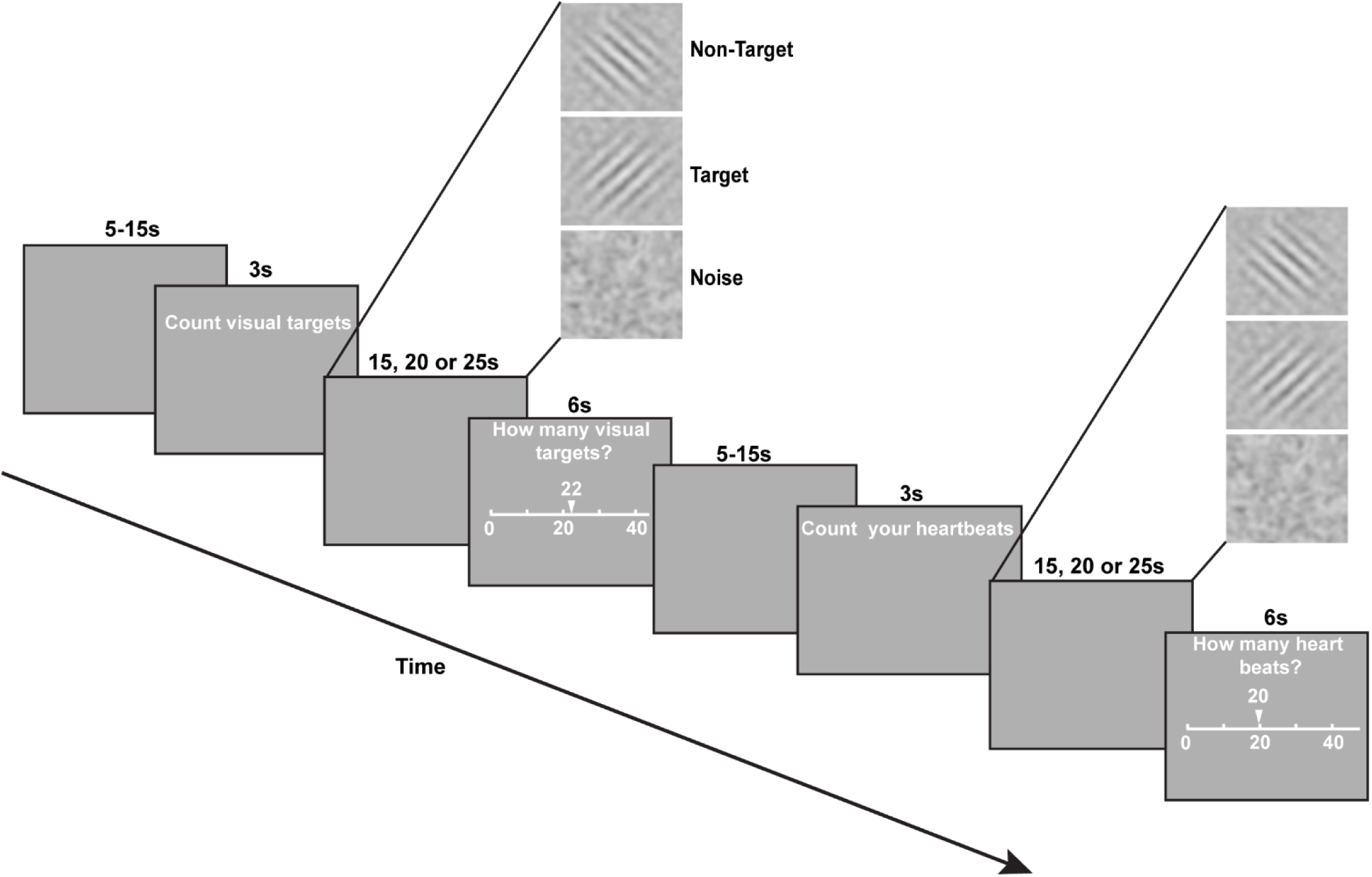
Experimental paradigm. The experiment employed a block design comprising exteroceptive (Visual Target Counting, VTC) and interoceptive (Heartbeat Counting, HBC) attention conditions. Each trial began with a 3 s instruction cue (“Count visual targets” or “Count your heartbeats”), followed by an active task block lasting 15, 20 or 25 s in pseudorandom order. During the VTC condition, participants counted centrally presented target Gabor patches (45° orientation) embedded within a continuous 10 Hz visual stream comprising distractor Gabor patches (−45° orientation) and visual noise. During the HBC condition, participants ignored the visual targets and silently counted their perceived heartbeats while maintaining visual fixation. Crucially, the identical 10 Hz visual stream remained present throughout both conditions, ensuring that bottom-up visual input was matched across tasks while attention was selectively directed towards either external visual stimuli or internal cardiac signals. Following each task block, participants reported the total number of visual targets or heartbeats counted during a 6 s response period by adjusting a slider displaying the selected numerical value using an MRI-compatible handheld button box. Consecutive task blocks were separated by a variable inter-trial interval (ITI) of 5–15 s, during which a blank screen was presented.

During VTC blocks, participants counted the number of rightward-tilted (45°) Gabor targets embedded within the continuous 10 Hz stream of visual noise and leftward-tilted distractors. The position of targets within the visual stream was yoked to previously collected cardiac data from an independent group of participants. Five target-timing sequences were used, corresponding to mean heart rates of 59, 66, 73, 74, and 76 beats per minute, with one sequence randomly selected for each participant. Thus, the temporal frequency and rhythmicity of the visual targets approximated those of naturally occurring heartbeats, reducing the possibility that participants could estimate target counts based simply on differences in temporal regularity between the two tasks. Distractors were inserted randomly according to a Poisson distribution, with a probability of 5% per stimulus in the 10Hz stream. During HBC blocks, participants silently counted their own perceived heartbeats while keeping their eyes open. Critically, the 10 Hz visual stimulus stream remained continuously present throughout both tasks. Thus, the physical visual input was identical during HBC and VTC (just as cardiac impulses continued during the visual task), and the only experimental manipulation was the instructed focus of attention.

Task blocks were separated by inter-trial rest periods lasting between 5 and 15 s (pseudorandomly determined), during which the visual stimulus stream was removed and participants viewed a blank screen before the next instruction cue. Following each task block, participants were given 6 s to report the total number of visual targets or heartbeats counted using an MRI-compatible button box. At the end of the scanning session, participants rated their overall confidence in their performance on a 6-point Likert scale (1 = not confident at all; 6 = extremely confident).

Throughout the entire scanning session, cardiac activity was continuously recorded using an infrared pulse oximeter attached to the index finger of the non-dominant hand. The visual task was piloted to achieve behavioural performance comparable to that of heartbeat counting, allowing differences in neural activity between conditions to be attributed primarily to the direction of attention rather than differences in task difficulty or sensory stimulation.

### MRI Data Acquisition

MRI data were acquired on a Siemens 3 T whole-body MRI scanner (Siemens Healthcare, Erlangen, Germany) at the Wellcome Centre for Human Neuroimaging, University College London, using a 12-channel head coil for functional imaging. Head motion was minimized by providing participants with instructions to remain still and by stabilizing the head with foam padding.

Functional images were acquired continuously using a T2*-weighted gradient-echo echo-planar imaging (EPI) sequence sensitive to blood oxygenation level-dependent (BOLD) contrast (TR = 3.36 s, TE = 30 ms, voxel size = 3 × 3 × 3 mm, matrix size = 64 × 64, 48 axial slices covering the whole brain, echo spacing = 0.5 ms, slice orientation = 30° relative to the anterior commissure–posterior commissure (AC–PC) plane).

At the end of the functional scanning session, a high-resolution T1-weighted structural image (1 mm isotropic resolution, TR = 18.7 ms, TE = 4.7 ms, flip angle = 20°, matrix size = 240 × 256 × 176) was acquired as part of the Multi-Parameter Mapping (MPM) protocol (Weiskopf et al., 2013) using a 32-channel head coil.

### fMRI Preprocessing

Functional and structural MRI data were preprocessed using the CONN functional connectivity toolbox (Whitfield-Gabrieli and Nieto-Castanon, 2012), which implements preprocessing routines from Statistical Parametric Mapping (SPM12; Wellcome Centre for Human Neuroimaging, London, UK).

Functional images were first corrected for head motion by rigid-body realignment and unwarping to account for susceptibility-by-motion interactions, followed by slice-timing correction. Structural T1-weighted images were co-registered to the mean functional image and segmented into grey matter, white matter, and cerebrospinal fluid (CSF). Structural and functional images were subsequently normalized to the Montreal Neurological Institute (MNI) template space and re-sampled to 2 x 2 x 2 mm isotropic voxels, followed by spatially smoothing with a 6 mm full width at half-maximum (FWHM) Gaussian kernel.

Motion-contaminated volumes were identified using the Artifact Detection Tools (ART) implemented within CONN, and scan-specific scrubbing regressors were generated to be included in the general linear modelling analysis as nuisance regressors.

### Statistical Analysis

Statistical analyses were performed using the General Linear Model (GLM) implemented in SPM12 (Friston et al., 1994). For each participant, the first-level design matrix comprised three regressors corresponding to the heartbeat counting task, the visual target counting task, and the rating period. Each task regressor was modelled as a boxcar function spanning the onset and duration of the corresponding block and convolved with the canonical hemodynamic response function.

To minimize the influence of motion-and physiology-related artefacts, several nuisance regressors were included in the first-level model. These comprised the six rigid-body head-motion parameters together with their temporal derivatives, scan-specific scrubbing regressors identified using the ART toolbox, five principal components extracted from white matter and five from cerebrospinal fluid using the anatomical CompCor (aCompCor) method (Behzadi et al., 2007), and physiological regressors generated using the TAPAS PhysIO toolbox (Kasper et al., 2017). The physiological model included six cardiac RETROICOR regressors together with their temporal derivatives, as well as a heart rate variability (HRV) regressor and its temporal derivative. A high-pass filter with a cut-off period of 128 s was applied to remove low-frequency signal drifts.

Following model estimation, first-level contrast images were generated for heartbeat counting relative to the implicit baseline (HBC > Rest baseline) and visual target counting relative to the implicit baseline (VTC > Rest baseline). These contrast images were entered into a second-level random-effects paired *t*-test, with participant modelled as a within-subject factor, to assess differential activity between conditions. This yielded the principal contrasts HBC > VTC and VTC > HBC. Statistical maps were thresholded at peak threshold of p = 0.001 followed by cluster-level false discovery rate (FDR) correction for multiple comparisons (*p*FDR < 0.05).

To facilitate interpretation of these differential contrasts, contrast estimates were extracted from the first-level HBC > Rest and VTC > Rest contrast images (SPM *con* images) within clusters identified by the second-level HBC > VTC and VTC > HBC analyses. These contrast estimates were used to determine whether significant between-condition differences reflected increased activation or decreased suppression, relative to the implicit baseline.

## Results

### 1. Behavioral performance was matched between the task conditions

Participants performed the heartbeat counting and visual target counting tasks with comparable levels of accuracy (Figure 2A). Mean accuracy for the HBC task was 81.9% (SD = 17.2; median = 89.3%) and 84% (SD = 11.8; median = 86.6%) for the VTC task. A Wilcoxon signed-rank test revealed no significant difference in accuracy between the two tasks (*p* = 0.6542).

**Figure 2.**
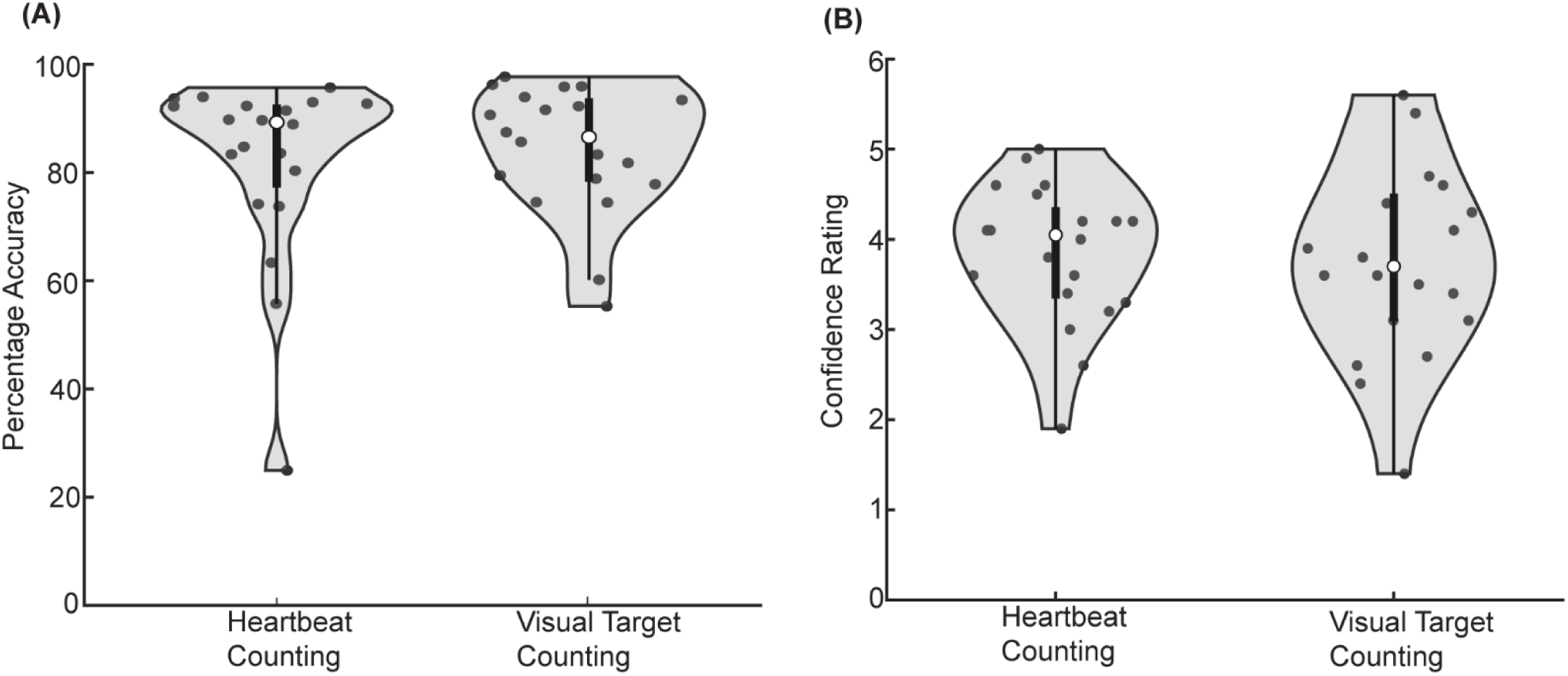
Behavioural performance during the heartbeat counting and visual target counting tasks. (A) Accuracy (%) during the HBC and VTC tasks. Accuracy did not differ significantly between the two conditions (Wilcoxon signed-rank test, *p* = 0.65), indicating comparable behavioural performance across tasks. (B) Confidence ratings following the HBC and VTC tasks. Confidence ratings were also comparable between the two conditions (Wilcoxon signed-rank test, *p* = 0.77). Violin plots illustrate the distribution of the data, boxplots indicate the median and interquartile range, and circles represent individual participants.

Self-rated confidence ratings were similarly matched across the two tasks (Figure 2B). Participants reported a mean confidence rating of 3.84 (SD = 0.8; median = 4.1) for the HBC and 3.8 (SD = 1.1; median = 3.7) for VTC. Confidence ratings did not differ significantly between the two task conditions (Wilcoxon signed-rank test, *p* = 0.7650).

The temporal frequencies of the five visual target sequences (59–76 targets/min) fell within the range of heart rates observed across participants during the experiment (55.7– 82.0 beats/min), confirming that the visual target frequencies were comparable to the cardiac frequencies encountered during heartbeat counting.

We additionally examined the correspondence between task accuracy and subjective confidence. Heartbeat-counting accuracy was positively associated with heartbeat-counting confidence (Spearman’s ρ = 0.50, p = 0.026), and a similar positive association was observed between visual target-counting accuracy and visual target-counting confidence (Spearman’s ρ = 0.47, p = 0.036). Thus, across participants, subjective confidence tracked objective performance in both the interoceptive and exteroceptive attention tasks.

Together, these behavioural findings indicate that the two tasks were comparable in overall accuracy and subjective confidence, while also demonstrating a positive correspondence between objective performance and subjective confidence within both tasks.

### 2. Visual target counting preferentially recruits visual areas and dorsal attention network

#### 2.1 Visual target counting enhanced activity within the foveal representation of the primary visual cortex

To identify regions preferentially engaged during externally directed attention, whole-brain activation during VTC was contrasted with that during HBC. This analysis revealed significantly greater activation within the bilateral occipital pole which corresponds to the foveal representation of the primary visual cortex (Figure 3i). Peak activations were observed in the left occipital pole (MNI = −18, −96, −10; *t*(19) = 7.92) and right occipital pole (MNI = 26, −96, −6; *t*(19) = 5.59), with both clusters surviving whole-brain false discovery rate correction (*pFDR* < 0.05).

**Figure 3.**
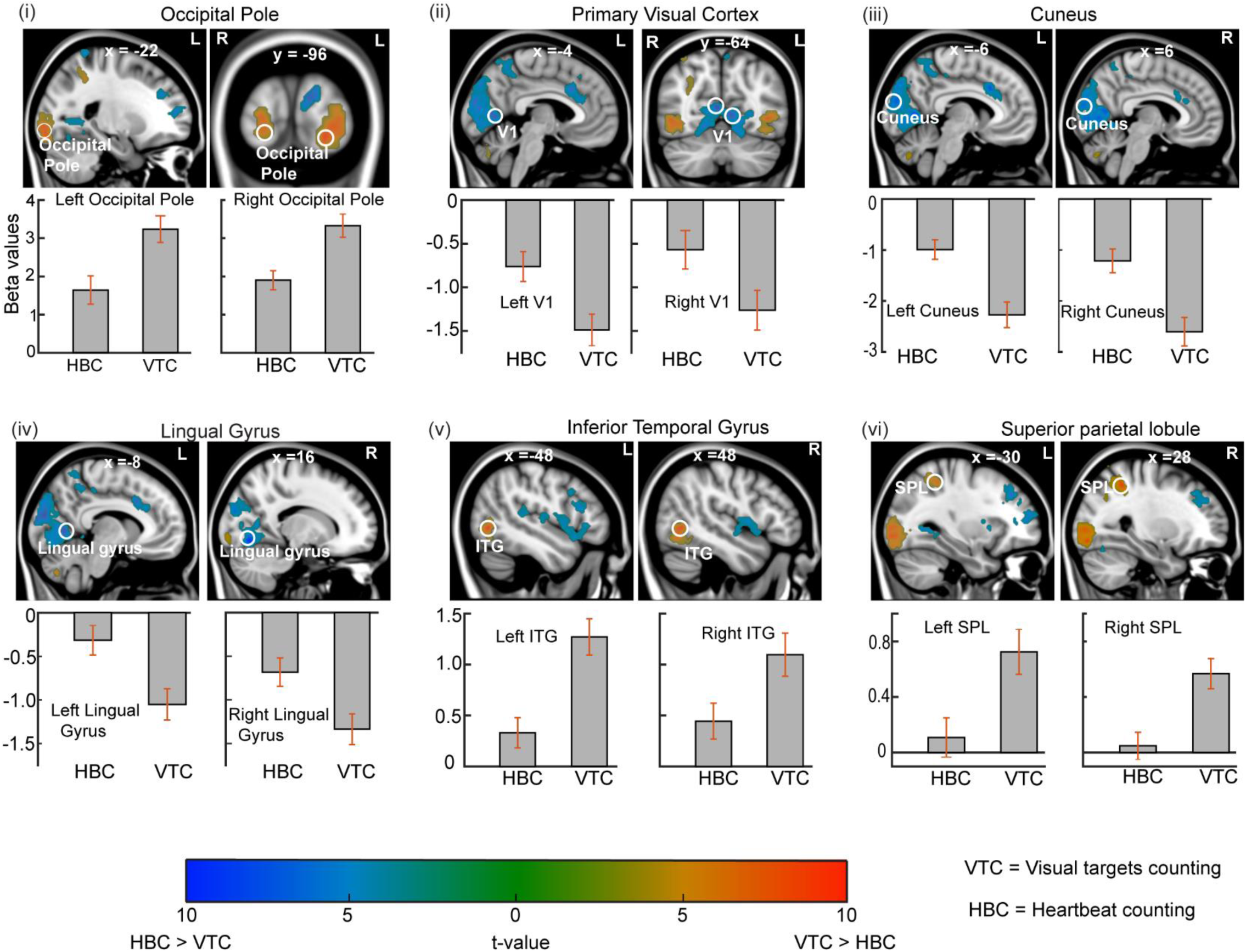
Visual exteroceptive attention differentially modulates visual cortex and recruits the dorsal attention network. Statistical parametric maps (voxel level threshold p <0.001 uncorrected; cluster-level FDR-corrected *p* < 0.05; displayed on MNI152 anatomical template) show whole-brain differences in activation between the Visual Target Counting (VTC) and Heartbeat Counting tasks (HBC). Orange–yellow clusters indicate regions exhibiting greater activation during VTC (VTC > HBC), whereas blue clusters indicate regions exhibiting greater activation during HBC (HBC > VTC) (colour bar: *t*-value). Circles indicate the peak voxels from which parameter estimates were extracted. Bar graphs beneath each panel show the mean parameter estimates (β values), relative to the inter-trial baseline, for the HBC and VTC tasks; error bars represent ±1 SEM. (i) Bilateral occipital pole (consistent with central visual-field representations). (ii) Medial occipital cortex (consistent with more peripheral visual representations) (iii) Cuneus. (iv) Lingual gyrus. (v) Inferior temporal gyrus (ITG). (vi) Superior parietal lobule (SPL). Slice coordinates are given in the (MNI) space (mm).

During the VTC task, participants continuously attended a small centrally presented region of the visual display while ignoring the surrounding visual field. Consistent with the retinotopic organization of the primary visual cortex, this attentional focus was therefore expected to preferentially engage the cortical representation of the fovea. To determine whether the greater occipital pole activation reflected enhancement during VTC or suppression during HBC, parameter estimates were extracted from each significant cluster. Both VTC and HBC elicited significant positive BOLD responses in foveal cortical areas in calcarine cortex near occipital pole relative to baseline, consistent with the continuous presentation of identical visual stimulation during both task conditions. However, activation was significantly greater during VTC than HBC, indicating that directing attention towards centrally presented visual targets selectively enhanced visually evoked responses within the cortical representation of the attended region of visual space.

#### 2.2 Visual target counting was associated with suppression of cortical representations of peripheral vision

In contrast to the enhancement observed within the foveal representation of primary visual cortex, parameter estimates revealed widespread suppression of activity relative to the resting baseline within the bilateral peripheral representations of primary visual cortex (anterior to the foveal representation), the cuneus, and the lingual gyrus (Figure 3ii–iv).

Consistent with these observations, the contrast (HBC > VTC) revealed significantly greater activity within the bilateral primary visual cortex (left peak: MNI = −4, −70, 8; *t*(19) = 6.74; right peak: MNI = 4, −64, 10; *t*(19) = 7.76), cuneus (left peak: MNI = −6, −88, 20; *t*(19) = 7.10; right peak: MNI = 6, −90, 16; *t*(19) = 6.35), and lingual gyrus (left peak: MNI = −8, −66, 2; *t*(19) = 6.85; right peak: MNI = 16, −70, −8; *t*(19) = 8.90), with all clusters surviving whole-brain FDR correction (*pFDR* < 0.05).

The extracted parameter estimates showed that the greater activity observed during HBC primarily reflected stronger suppression during VTC rather than enhanced activation during HBC. Activity within these regions remained below baseline during both tasks, but suppression was significantly attenuated during HBC. Together with the enhanced activation observed in the occipital pole, these findings indicate that externally directed visual attention was accompanied by a reciprocal pattern of facilitation and suppression across the central and peripheral representations of early visual cortex.

#### 2.3 Higher-order visual cortex exhibited stronger activation during visual target counting

Beyond early visual cortex, VTC also elicited significantly greater activation within higher-order visual cortex, including the bilateral inferior temporal gyrus (Figure 3v). Peak activations were identified in the left inferior temporal gyrus (MNI = −48, −68, −2; *t*(19) = 8.81) and right inferior temporal gyrus (MNI = 48, −60, −2; *t*(19) = 7.37), with both clusters surviving whole-brain FDR correction (*pFDR* < 0.05).

The extracted parameter estimates demonstrated significant positive BOLD responses relative to baseline during both task conditions, with significantly greater activation during VTC than HBC.

#### 2.4 Visual target counting recruited the dorsal attention network

The VTC > HBC contrast further revealed robust bilateral activation within the superior parietal lobule (SPL), a principal node of the dorsal frontoparietal attention network (Figure 3vi). Significant activation was observed in the left SPL (MNI = −30, −44, 50; *t*(19) = 6.01) and right SPL (MNI = 28, −46, 48; *t*(19) = 8.51), with both clusters surviving whole-brain FDR correction (*pFDR* < 0.05).

The extracted parameter estimates showed significant positive BOLD responses relative to the resting baseline during VTC, whereas activity during HBC did not differ significantly from baseline (left SPL: *t*(19) = 0.75, *p* = 0.47; right SPL: *t*(19) = 0.49, *p* = 0.64). These findings indicate that recruitment of the SPL was selective for externally directed visual attention.

At a more liberal statistical threshold (*p* < 0.001, uncorrected), greater activation during VTC was also observed within the bilateral frontal eye fields (FEF) (left peak: MNI = −30, −8, 50; *t*(19) = 3.04; right peak: MNI = 32, −8, 52; *t*(19) = 3.44) which is another important node of the dorsal attention network. Together with the SPL and FEF activation showed that the visual attention engaged the canonical dorsal attention network.

### 3. Brain regions recruited during interoceptive attention

#### 3.1 Heartbeat counting recruited the posterior insula

Whole-brain comparison of the HBC with VTC (HBC > VTC) showed robust bilateral activation of the posterior insula (Figure 4i). In the left hemisphere, two distinct local maxima were identified within the posterior insula: a posterior peak (MNI = −38, −16, 2; *t*(19) = 5.16) and a more anterior peak (MNI = −38, −2, −4; *t*(19) = 5.13). In the right hemisphere, a single local maximum was observed at MNI = (42, −8, −8; *t*(19) = 6.67). All of these clusters survived the whole-brain FDR correction (*pFDR* < 0.05). The two left posterior insular local maxima are labelled “p” (posterior) and “a” (anterior) in Figure 4(i).

**Figure 4.**
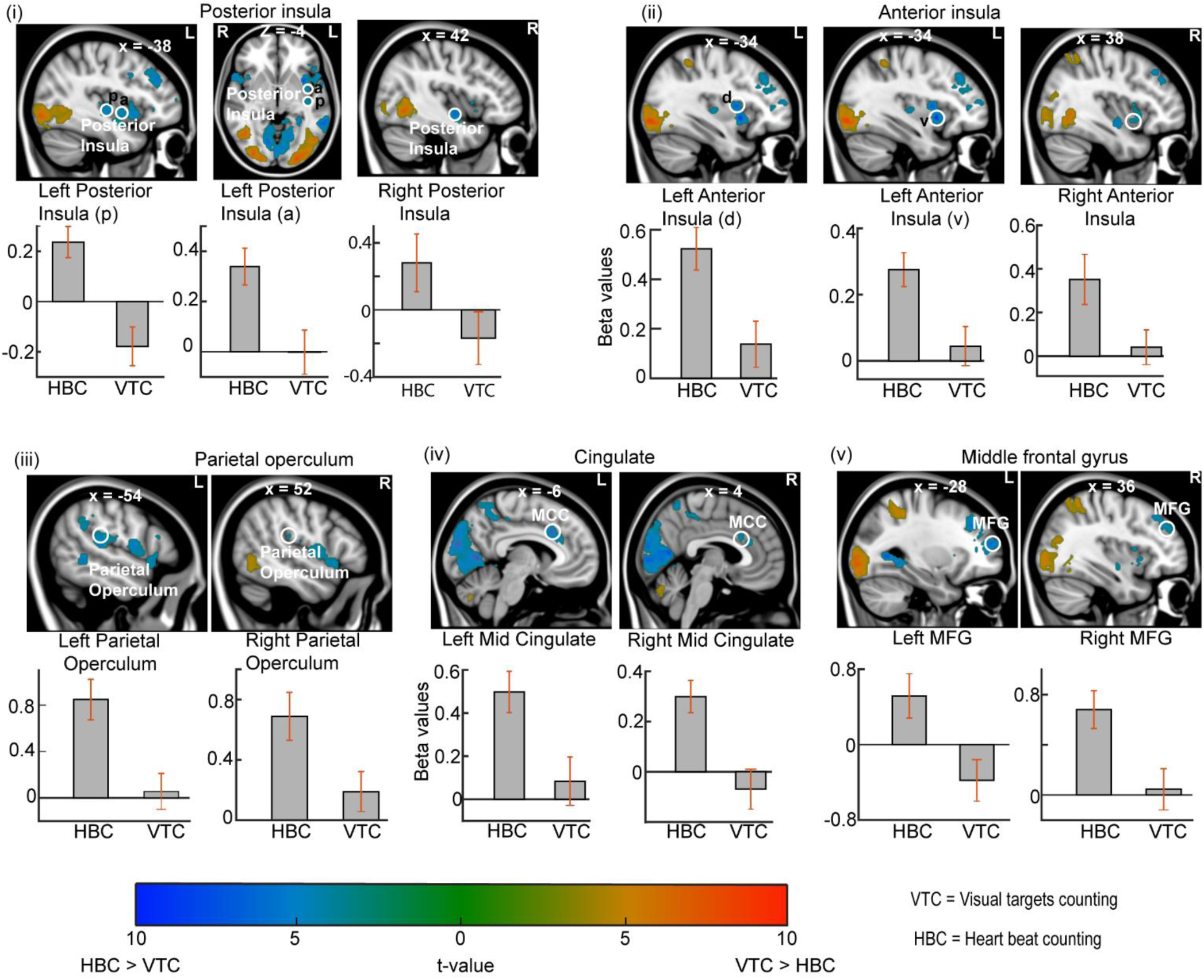
Cardiac interoceptive attention recruits a distributed cortical network. Statistical parametric maps (display as in figure 3) show whole-brain differences in activation between the Heartbeat Counting (HBC) and Visual Target Counting (VTC) tasks. Orange–yellow clusters indicate regions exhibiting greater activation during HBC than VTC (colour bar: *t*-value). Circles indicate the peak voxels from which parameter estimates were extracted. Bar graphs beneath each panel show the mean parameter estimates (β values), relative to the inter-trial baseline, for the HBC and VTC tasks; error bars represent ±1 SEM. (i) Posterior insula, exhibiting greater activation during HBC. Two local maxima are identified in the left hemisphere (posterior, p; anterior, a). (ii) Anterior insula, exhibiting preferential activation during HBC. Two local maxima are identified in the left hemisphere (dorsal, d; ventral, v). (iii) Bilateral parietal operculum (secondary somatosensory cortex, SII), exhibiting greater activation during HBC. (iv) Bilateral anterior/mid-cingulate cortex, exhibiting greater activation during HBC. (v) Bilateral middle frontal gyrus, exhibiting greater activation during HBC. Slice coordinates are given in the MNI space (mm). L, left hemisphere; R, right hemisphere.

The extracted parameter estimates showed that all three posterior insular loci exhibited significant positive BOLD responses during HBC relative to the resting baseline. During VTC, however, only the posterior (“p”) peak within the left posterior insula showed a significant negative BOLD response relative to baseline (*t*(19) = −2.25, *p* < 0.05). In contrast, activity at the more anterior left posterior-insular (“a”) peak (*t*(19) = −0.015, *p* = 0.98) and the right posterior insular peak (*t*(19) = −1.05, *p* = 0.31) did not differ significantly from baseline. Thus, heartbeat counting robustly recruited the posterior insula bilaterally, whereas suppression during the visual attention task was confined to the most posterior subdivision of the left posterior insula.

#### 3.2 Heartbeat counting recruited the anterior insula

The HBC > VTC contrast showed robust bilateral activation of the anterior insula (Figure 4ii). In the left hemisphere, two distinct local maxima were identified within the anterior insula: a dorsal peak (MNI = −34, 6, 8; *t*(19) = 7.28) and a ventral peak (MNI = −34, 8, −4; *t*(19) = 9.34). The clusters are marked “d” (dorsal) and “v” (ventral) in Figure 4(ii). In the right hemisphere, a single local maximum was observed at MNI = (38, 8, −6; *t*(19) = 3.90).

The extracted parameter estimates showed that all anterior insular loci exhibited significant positive BOLD responses during HBC relative to the resting baseline. In contrast, none of the anterior insular loci showed significant activation during VTC. Activity at the left dorsal peak did not differ significantly from baseline (*t*(19) = 1.43, *p* = 0.17), nor did activity at the left ventral peak (*t*(19) = 0.73, *p* = 0.47) or the right anterior insular peak (*t*(19) = 0.50, *p* = 0.63). Thus, heartbeat counting robustly recruited both dorsal and ventral subdivisions of the anterior insula bilaterally, whereas visual target counting did not evoke significant activation within these regions.

#### 3.3 Heartbeat counting additionally recruited the parietal operculum

Whole-brain comparison of the Heartbeat Counting and Visual Target Counting tasks (HBC > VTC) also revealed robust bilateral activation of the parietal operculum (secondary somatosensory cortex; Figure 4iii). In the left hemisphere, a local maximum was identified at (MNI = −54, −32, 18; *t*(19) = 5.05), while a homologous activation was observed in the right hemisphere at (MNI = 52, −28, 20; *t*(19) = 4.17). Both clusters survived whole-brain FDR correction (pFDR < 0.05). Inspection of the extracted parameter estimates demonstrated significant positive BOLD responses during HBC relative to the resting baseline in both hemispheres, whereas activity during VTC remained close to baseline.

#### 3.4 Heartbeat counting recruited a fronto-cingulate network

Whole-brain comparison of the Heartbeat Counting and Visual Target Counting tasks (HBC > VTC) additionally revealed robust activation within a fronto-cingulate network comprising the anterior/mid-cingulate cortex and the middle frontal gyrus (Figure 4iv,v). Bilateral activation was observed within the anterior/mid-cingulate cortex, with local maxima at (MNI = −6, 18, 34; *t*(19) = 7.00) in the left hemisphere and (MNI =4, 20, 26; *t*(19) = 4.43) in the right hemisphere. The HBC > VTC contrast also revealed significant bilateral activation of the middle frontal gyrus, with peaks at (MNI = −28, 54, 16; *t*(19) = 6.29, *p* < 0.001) on the left and (MNI =36, 38, 36; *t*(19) = 5.40) on the right. All clusters survived whole-brain FDR correction at p < 0.05.

The extracted parameter estimates demonstrated positive BOLD responses during HBC relative to the resting baseline across both the anterior/mid-cingulate cortex and the middle frontal gyrus, whereas activity during VTC remained close to baseline. Thus, heartbeat counting engaged a distributed fronto-cingulate network that was not significantly recruited during visual target counting.

### 4. Across task suppression between exteroceptive and interoceptive attention

To systematically evaluate which brain regions recruited during interoceptive attention were selectively suppressed during exteroceptive attention, we inclusively masked the whole-brain HBC > VTC contrast with voxels exhibiting significant deactivation during the Visual Target Counting (VTC) task relative to the resting baseline (VTC < baseline). That is, those voxels which were suppressed during VTC but were released from suppression during HBC. Consistent with the observations described in Section 3.1 (Figure 4i), this analysis identified the most posterior subdivision of the left posterior insula, indicating that the greater HBC > VTC response within this primary interoceptive region was attributable, in part, to suppression during visual attention. Beyond the posterior insula, the analysis identified a restricted set of higher-order cortical regions, including the bilateral anterior precuneus, a medial frontal region encompassing the anterior cingulate cortex, and the left middle frontal gyrus (Figure 5). Extracted parameter estimates showed that the greater HBC > VTC responses in these brain regions arose primarily from suppression during exteroceptive attention rather than enhanced recruitment during interoceptive attention.

**Figure 5.**
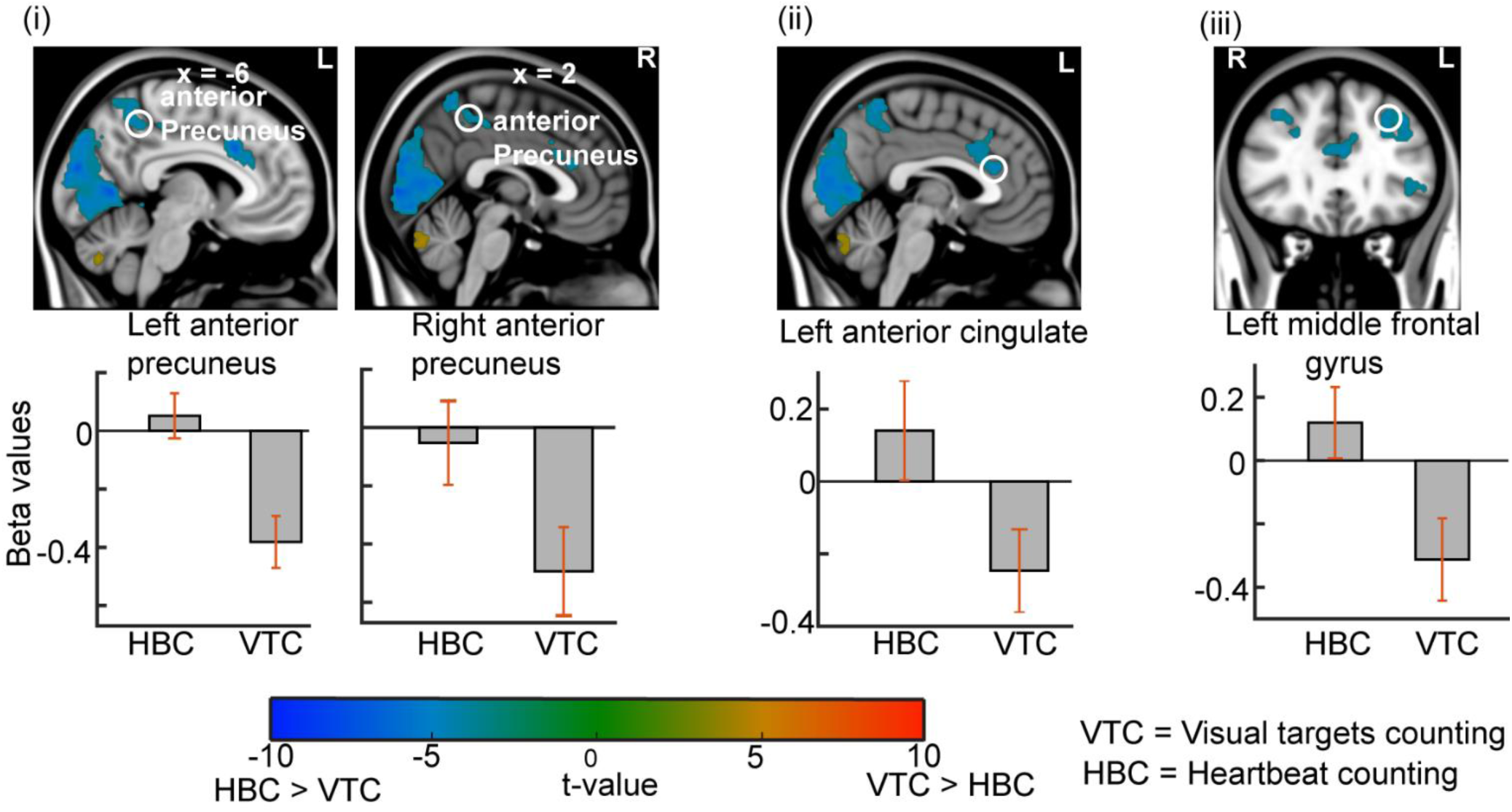
Selective suppression during visual exteroceptive attention. Statistical parametric maps show the whole-brain contrast Heartbeat Counting (HBC) > Visual Target Counting (VTC), inclusively masked by voxels exhibiting significant deactivation during the VTC task relative to the inter-trial baseline (VTC < baseline). This analysis identified regions in which the HBC > VTC difference was driven by selective suppression during visual exteroceptive attention rather than enhanced activation during cardiac interoceptive attention. Circles indicate the peak voxels from which parameter estimates were extracted. Bar graphs beneath each panel show the mean parameter estimates (β values), relative to the inter-trial baseline, for the HBC and VTC tasks; error bars represent ±1 SEM. (i) Bilateral anterior precuneus, exhibiting suppression below baseline during VTC while remaining close to baseline during HBC. (ii) Anterior cingulate cortex, exhibiting the same pattern of selective suppression during VTC. (iii) Left middle frontal gyrus, exhibiting suppression during VTC while remaining close to baseline during HBC. Slice coordinates are given in the MNI space (mm). L, left hemisphere; R, right hemisphere.

By contrast, the reciprocal masking analysis, in which the VTC > HBC contrast was inclusively masked with voxels exhibiting significant deactivation during heartbeat counting relative to the resting baseline (HBC < baseline), revealed no suprathreshold clusters. Thus, no brain regions preferentially recruited during visual attention showed evidence of selective suppression during interoceptive attention.

### 5. Correlation between behavioral data and BOLD activity

To identify brain regions associated with individual differences in task performance, we performed a whole-brain multiple regression analysis at the second level. Four behavioral measures were entered simultaneously into the design matrix: (i) heartbeat counting accuracy, (ii) visual target counting accuracy, (iii) confidence ratings for the HBC, and (iv) confidence ratings for the VTC. This analysis therefore identified brain regions whose activity was uniquely associated with each behavioral measure after accounting for the shared variance with the remaining behavioral variables.

A significant negative association with heartbeat counting accuracy was observed within the right inferior parietal lobule (Figure 6; peak MNI = 46, −50, 54; t(15) = 5.48), after whole-brain FDR correction (*pFDR* < 0.05). That is, participants with higher heartbeat counting accuracy exhibited smaller HBC > VTC contrast estimates within this region. No significant whole-brain corrected correlations were observed for visual target counting accuracy, heartbeat counting confidence, or visual target counting confidence following correction for multiple comparisons. At an uncorrected threshold of p < 0.001, exploratory analyses revealed positive correlations between heartbeat-counting confidence ratings and HBC > VTC contrast estimates in the left (MNI = −22, 48, −12; t(15) = 4.86) and right (MNI = 34, 44, −10; t(15) = 4.29) anterior prefrontal cortex. Although these effects did not survive correction for multiple comparisons, their localisation to anterior prefrontal cortex is notable given the involvement of this region in confidence and metacognitive judgements reported in previous studies (Fleming et al., 2010; Fleming and Dolan, 2012).

**Figure 6.**
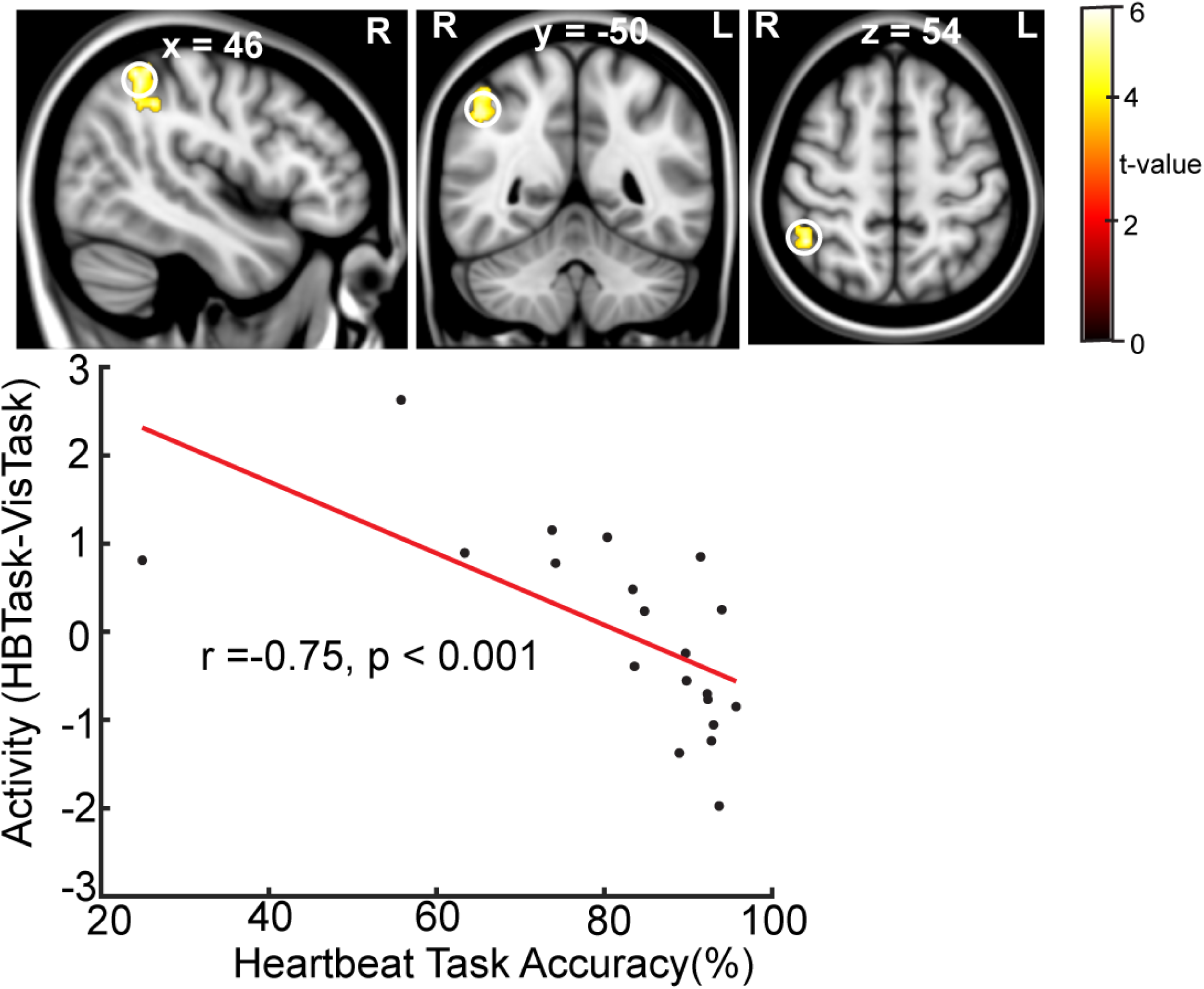
Brain–behaviour relationship for cardiac interoceptive accuracy. Statistical parametric maps show the results of a whole-brain multiple regression analysis relating regional BOLD activity to behavioural performance. Four behavioural measures were entered simultaneously into the second-level design matrix: heartbeat counting accuracy, visual target counting accuracy, heartbeat counting confidence, and visual target counting confidence. After partialling out the shared variance associated with the remaining behavioural measures, a significant negative association between heartbeat counting accuracy and BOLD activity was observed in the right angular inferior parietal lobule. The scatter plot illustrates the adjusted relationship between heartbeat counting accuracy and the extracted HBC > VTC contrast estimates from the peak voxel, after accounting for the remaining behavioural regressors. Higher heartbeat counting accuracy was associated with smaller HBC > VTC contrast estimates. Slice coordinates are given in MNI space (mm). R, right hemisphere.

To facilitate comparison with previous studies, e.g. (Critchley et al., 2004), we also performed secondary analyses in which each behavioral measure was entered individually as a single regressor. Unlike the multiple regression analysis, these analyses do not account for the shared variance between behavioral measures and should therefore be considered complementary. When heartbeat counting accuracy was considered alone, significant positive correlations were observed bilaterally within the thalamus in the left (peak MNI = −14, −10, 14; t(19) = 5.27) and right (peak MNI =-2,-12, 12; t(19) =4.88), both clusters surviving whole-brain FDR correction (*pFDR* < 0.05). A positive correlation was also observed in the left anterior insula (peak MNI = −38, 10, −8; t(19) = 4.20), although this effect was evident only at an uncorrected threshold of (p < 0.001). Heartbeat counting confidence ratings were positively associated with bilateral anterior orbital gyri (left peak MNI = −30, 48, −12; t(19) = 4.49; right peak MNI = 34, 44,−12; t(19) = 4.69), although these effects were likewise observed only at an uncorrected threshold (p < 0.001) and did not survive correction for multiple comparisons. No significant correlations with either visual target counting accuracy or visual target counting confidence were observed.

Peak MNI coordinates for the contrasts HBC>VTC and VTC>HBC are given in Tables 1 and 2 respectively.

**Table 1:** Brain regions showing greater activity during heartbeat counting than visual target counting (HBC > VTC). Clusters surviving whole-brain cluster-level FDR correction (*p*FDR < 0.05) at a voxel-forming threshold of *p* < 0.001 (uncorrected) are reported. MNI coordinates and *t*-values correspond to local maxima within each significant cluster.

| Brain region | Peak MNI coordinates | Peak $t$-value | Cluster size (Number of voxels) |
| --- | --- | --- | --- |
| Left Anterior Insula | -34, 8, -4 | 9.34 | 982 |
| Left Anterior Insula | -34, 6, 8 | 7.28 |  |
| Left Inferior Frontal Gyrus | -50, 26, 0 | 5.41 |  |
| Right Lingual Gyrus | 16, -70, -8 | 8.90 | 5076 |
| Right Calcarine Cortex | 4, -64, 10 | 7.76 |  |
| Left Middle Temporal Gyrus | -62, -46, -2 | 8.09 | 335 |
| Left Middle Temporal Gyrus | -62, -42, -10 | 5.26 |  |
| Left Middle Temporal Gyrus | -62, -56, -4 | 4.88 |  |
| Left Middle Cingulate | -6, 18, 34 | 7.00 | 411 |
| Left Anterior Cingulate | -4, 28, 22 | 5.13 |  |
| Left Medial Superior Frontal Gyrus | -10, 28, 28 | 5.01 |  |
| Right Posterior Insula | 42, -8, -8 | 6.67 | 903 |
| Right Central Operculum | 56, 8, 2 | 6.45 |  |
| Right Temporal Pole | 54, 16, -6 | 5.73 |  |
| Left Precuneus | -10, -50, 60 | 6.64 | 485 |
| Left Superior Parietal Lobule | -22, -48, 72 | 5.35 |  |
| Left Precuneus | -6, -46, 48 | 5.16 |  |
| Left Medial Precentral Gyrus | -16, -32, 44 | 6.34 | 97 |
| Left Middle Frontal Gyrus | -28, 54, 16 | 6.29 | 271 |
| Left Middle Frontal Gyrus | -36, 56, 12 | 4.83 |  |
| Left Middle Frontal Gyrus | -28, 34, 36 | 6.03 | 506 |
| Left Middle Frontal Gyrus | -38, 36, 32 | 5.80 |  |
| Superior Frontal Gyrus | -16, -2, 72 | 5.95 | 163 |
| Left Parietal Operculum | -60, -28, 16 | 5.94 | 990 |
| Left Supramarginal Gyrus | -66, 28, 28 | 5.93 |  |
| Left Supramarginal Gyrus | -66, -36, 24 | 5.67 |  |
| Right Middle Frontal Gyrus | 38, 32, 42 | 5.61 | 319 |
| Right Middle Frontal Gyrus | 36, 38, 36 | 5.40 |  |
| Right Middle Frontal Gyrus | 28, 40, 32 | 4.96 |  |
| Left Middle Frontal Gyrus | -44, 10, 34 | 5.34 | 184 |
| Left Posterior Insula | -38, -16, 2 | 5.16 | 150 |
| Left Transverse Temporal Gyrus | -50, -16, 8 | 4.30 |  |
| Right Supramarginal Gyrus | 56, -40, 30 | 4.73 | 78 |
| Right Parietal Operculum | 52, -28, 20 | 4.17 | 92 |
| Right Parietal Operculum | 60, -24, 24 | 3.94 |  |
| Left Middle Cingulate | -6, 0, 42 | 4.54 | 55 |
| Left Inferior Frontal Gyrus | -48, 40, 4 | 4.31 | 55 |
| Left Inferior Frontal Gyrus | -50, 34, 12 | 3.71 |  |

**Table 2.** Brain regions showing greater activity during visual target counting than heartbeat counting (VTC > HBC). Clusters surviving whole-brain cluster-level FDR correction (*p*FDR < 0.05) at a voxel-forming threshold of *p* < 0.001 (uncorrected) are reported. MNI coordinates and *t*-values indicate local maxima within each significant cluster.

| Brain region | MNI coordinates | t-value | Cluster Size<br>(Number of<br>voxels) |
| --- | --- | --- | --- |
| Left Inferior Occipital Gyrus | -48, -68, -2 | 8.81 | 1762 |
| Left Inferior Occipital Gyrus | -36, -86, -8 | 8.60 |  |
| Left Occipital Pole | -18, -96, -10 | 7.92 |  |
| Right Superior Parietal Lobule | 28, -46, 48 | 8.51 | 449 |
| Right Superior Parietal Lobule | 36, -52, 58 | 5.22 |  |
| Right Angular Gyrus | 36, -60, 56 | 4.54 |  |
| Right Occipital Fusiform Gyrus | 28, -84, -10 | 8.23 | 1006 |
| Right Inferior Occipital Gyrus | 32, -86, -2 | 7.39 |  |
| Right Inferior Temporal Gyrus | 46, -58, 0 | 6.94 | 491 |
| Left Superior Parietal Lobule | -30, -44, 50 | 6.01 | 277 |
| Left Cerebellum Exterior | -10, -74, -42 | 5.48 | 153 |
| Right Cerebellum | 2, -74, -26 | 4.63 |  |
| Right Cerebellum | 8, -74, -38 | 4.60 |  |

## Discussion

The present study investigated how the human brain allocates attention between competing internal bodily signals and external sensory information using a controlled comparison between cardiac interoceptive and visual exteroceptive attention. By maintaining identical visual stimulation during the active periods of both tasks while matching behavioral accuracy and subjective confidence, our experimental design isolated differences in attentional allocation from differences in sensory input and reduced the likelihood that they were driven by differences in task performance. Cardiac interoceptive and visual exteroceptive attention engaged largely distinct cortical networks. Whereas interoceptive attention recruited a distributed insula-centered network supporting interoceptive processing and cognitive control, exteroceptive attention engaged dorsal frontoparietal attentional regions together with selective modulation of visual cortex. Importantly, attentional allocation was also associated with suppression of activity in regions involved in processing the competing source of information. Together, these findings suggest that the allocation of attention between bodily and external information involves coordinated modulation of interoceptive, attentional and sensory systems.

Cardiac interoceptive attention recruited a distributed network encompassing the posterior and anterior insula, secondary somatosensory cortex, anterior cingulate cortex and middle frontal gyrus. This pattern is broadly consistent with previous neuroimaging studies identifying these regions as key components of the cortical network supporting the perception and cognitive evaluation of internal bodily signals (Critchley et al., 2004; Pollatos et al., 2007; Simmons et al., 2013; Wiebking et al., 2014b; Wiebking et al., 2014a; Tan et al., 2018; Failla et al., 2020). The posterior insula has long been regarded as the primary cortical recipient of ascending interoceptive afferents and is thought to provide a representation of the body’s physiological state, whereas the anterior insula has been implicated in integrating these bodily signals with cognitive and motivational processes that support conscious access to interoceptive sensations (Craig, 2002, 2009; Critchley and Harrison, 2013). Beyond the insula, activation of the secondary somatosensory cortex may reflect the contribution of somatic afferent signals accompanying each heartbeat, such as subtle chest wall or vascular sensations, which have been proposed to provide primary or additional information during heartbeat perception (Khalsa et al., 2009; Couto et al., 2014). The anterior cingulate cortex and middle frontal gyrus were preferentially recruited during cardiac interoceptive attention despite the two tasks being closely matched for behavioral performance and involving comparable counting and working memory requirements. This finding suggests that these regions contribute not simply to generic executive processes but to the additional top-down control required to sustain attention towards relatively weak internal bodily signals while resisting distraction from continuously available external sensory information.

Visual exteroceptive attention recruited bilateral superior parietal lobule together with selective modulation of early visual cortex, a pattern that closely accords with established models of visual selective attention. The superior parietal lobule is a key component of the dorsal frontoparietal attention network and has been consistently implicated in the voluntary allocation of attention towards behaviorally relevant visual information (Kastner and Ungerleider, 2000; Corbetta and Shulman, 2002). At the sensory level, directing attention towards centrally presented visual targets enhanced activity within the occipital pole, while suppressing activity within more medial occipital regions. This spatial pattern is consistent with the established eccentricity organisation of early visual cortex, in which central visual-field representations are concentrated around the occipital pole and progressively more peripheral representations extend anteriorly along medial occipital cortex (Wandell et al., 2005; Wandell et al., 2007). This pattern closely parallels previous behavioral, electrophysiological and neuroimaging studies demonstrating that visual attention enhances processing of behaviorally relevant sensory information while suppressing competing or behaviorally irrelevant inputs (Brefczynski and DeYoe, 1999; Somers et al., 1999; Carrasco, 2011).

A notable finding of the present study is that attentional allocation between internal bodily signals and external sensory information was associated not only with enhancement of task-relevant neural systems but also with selective suppression of regions associated with the competing source of information. Specifically, directing attention towards external visual stimuli suppressed activity within the posterior insula relative to the implicit resting baseline. The pattern within visual cortex was more complex. Medial occipital regions showed strong suppression relative to the implicit baseline during visual target counting. This suppression was significantly attenuated during heartbeat counting, consistent with the reduced requirement for spatially selective visual attention when participants redirected attention from the visual targets towards their heartbeats. Importantly, however, activity in these regions did not return to baseline during heartbeat counting but remained significantly below it. In contrast, the occipital pole remained positively activated during both tasks, although more strongly during visual target counting, consistent with the continuous presentation of the central visual stimulus. The continued suppression within medial occipital cortex during heartbeat counting has at least two possible interpretations. It may reflect persistence, albeit to a lesser degree, of the visual cortical suppression present during visual target counting. Alternatively, because the visual input remained continuously present but task-irrelevant during heartbeat counting, it may reflect active down-regulation of visual processing when attention is directed towards internal bodily signals.

The suppression observed during externally directed attention also extended beyond the posterior insula to the anterior precuneus, anterior cingulate cortex and left middle frontal gyrus. In these regions, the greater HBC relative to VTC activity primarily reflected suppression during visual target counting rather than increased recruitment during heartbeat counting. The involvement of the anterior precuneus is particularly notable given growing evidence for a role of this region in bodily representation. Lesion and direct cortical stimulation studies have implicated the anterior precuneus in bodily awareness and body schema, with stimulation producing alterations in the physical and spatial experience of the self (Herbet et al., 2019; Lyu et al., 2023). Importantly, anterior precuneus sites associated with these bodily experiences appear to be functionally distinct from adjacent posterior medial regions belonging to the default mode network (Lyu et al., 2023). Suppression of the anterior precuneus during visual target counting may therefore reflect reduced engagement of bodily self-related processing when attention is strongly directed towards external visual information.

An additional finding was that individual differences in cardiac interoceptive accuracy were associated with the differential recruitment of the right inferior parietal lobule (IPL). Greater heartbeat-counting accuracy was associated with smaller HBC > VTC contrast estimates within this region. The IPL has been implicated in attentional orienting towards behaviourally relevant events in the external environment (Corbetta and Shulman, 2002; Corbetta et al., 2008). Within the context of the present study, the reduced HBC relative to VTC response in individuals with higher interoceptive accuracy may therefore reflect reduced engagement of externally directed attentional processes while attention is focused on internal bodily signals. This finding suggests that successful cardiac interoception may depend not only on the processing of internal bodily signals but also on limiting the influence of neural systems supporting externally directed attention.

Accuracy and subjective confidence were positively associated across participants for both heartbeat counting and visual target counting, indicating that subjective confidence tracked objective performance across both interoceptive and exteroceptive attention. This correspondence between performance and confidence is relevant to metacognitive insight, which concerns the extent to which subjective judgements reflect objective performance. Exploratory neuroimaging analyses further showed that heartbeat-counting confidence was positively associated with activity in regions of the bilateral anterior prefrontal cortex which have previously been implicated in confidence and metacognitive judgements (Fleming et al., 2010; Fleming and Dolan, 2012). Although these effects did not survive correction for multiple comparisons and should therefore be interpreted cautiously, they raise the possibility that evaluation of confidence in cardiac interoceptive performance engages higher-order prefrontal systems involved in evaluating perceptual judgements.

Several limitations should be considered when interpreting the present findings. First, cardiac interoception was assessed using the heartbeat counting task, which, although widely used, reflects not only sensitivity to cardiac signals but also sustained attention, working memory and other cognitive processes (Desmedt et al., 2018; Desmedt et al., 2020; Ferentzi et al., 2022; Desmedt et al., 2023). However, because the visual target counting task was closely matched for counting, attentional demands and behavioral performance, these non-interoceptive components were minimized in the present comparison. Second, although the spatial pattern of visual cortical modulation was consistent with the established eccentricity organisation of early visual cortex, individual retinotopic mapping was not performed. The observed effects therefore cannot be definitively assigned to specific central or peripheral visual-field representations. Finally, the present study focused exclusively on cardiac interoception. Future studies should determine whether the allocation of attention between internal bodily signals and external sensory information observed here also characterizes other interoceptive modalities. Such findings would establish whether the principles identified in the present study reflect a general mechanism for balancing attention between the body and the external environment.

## Funding Acknowledgement

This work was supported by the Medical Research Council, UK (MR-T032553-1, awarded to T.D.G.).

